# OmniSegger: A time-lapse image analysis pipeline for bacterial cells

**DOI:** 10.1101/2024.11.25.625259

**Authors:** Teresa W. Lo, Kevin J. Cutler, H. James Choi, Paul A. Wiggins

## Abstract

Time-lapse microscopy is a powerful tool for studying the cell biology of bacterial cells. The development of pipelines that facilitate the automated analysis of these datasets is a long-standing goal of the field. In this paper, we describe *OmniSegger*, an updated version of our *SuperSegger* pipeline, developed as an open-source, modular, and holistic suite of algorithms whose input is raw microscopy images and whose output is a wide range of quantitative cellular analyses, including dynamical cell cytometry data and cellular visualizations. The updated version described in this paper introduces two principal refinements: (i) robustness to cell morphologies and (ii) support for a range of common imaging modalities. To demonstrate robustness to cell morphology, we present an analysis of the proliferation dynamics of *Escherichia coli* treated with a drug that induces filamentation. To demonstrate extended support for new image modalities, we analyze cells imaged by five distinct modalities: phase-contrast, two brightfield modalities, and cytoplasmic and membrane fluorescence. Together, this pipeline should greatly increase the scope of tractable analyses for bacterial microscopists.

## I. INTRODUCTION

Time-lapse microscopy is a powerful tool for understanding the structure and function of bacterial systems. In a single field of view, it is possible to capture the dynamics of hundreds to thousands of cells simultaneously [1]. The quantitative analysis of this image data can involve a wide range of cell cytometry, defined as the determination of morphological characteristics. Although these algorithms have a long history [2], the advent of machine-learning-based approaches has greatly facilitated the development of ever more robust and precise tools [3].

Ten years ago, we developed *SuperSegger*, an image analysis pipeline for the analysis of the time-dependent fluorescence localization in bacterial cells [4]. Its development was motivated by the challenge of performing a proteome-wide time-lapse analysis of localization of nearly all proteins with non-diffuse localization in *Escherichia coli* [1, 5]. However, since releasing the original SuperSegger analysis pipeline, we have discovered numerous scenarios in which the package fails to produce acceptable results.

In this paper, we highlight two types of analyses that will serve as the motivation for the development of a new package: diverse cell morphologies and the use of alternative image modalities. We will demonstrate a new image analysis pipeline, OmniSegger, which combines a new segmentation package we recently developed, Omnipose [3], with the existing analysis pipeline, SuperSegger [4]. We demonstrate that OmniSegger has superior performance to existing pipelines, both in the context of imaging standard time-lapse data, as well as vastly superior performance in the context of the morphology and modality challenges.

## II. RESULTS

### OmniSegger pipeline overview

The overall goal of the OmniSegger pipeline is to provide a open-source, modular, and holistic suite of algorithms whose input is raw multichannel microscopy images and whose output is a comprehensive range of quantitative cellular analyses presented in a user-friendly format that does not require coding expertise. However, we have attempted to make the algorithms modular to facilitate the use of individual modules by users who wish to code their own custom pipelines. A schematic representing the OmniSegger pipeline and a gallery of images and analyses generated using OmniSegger is shown in Fig. 1.

**FIG. 1.**
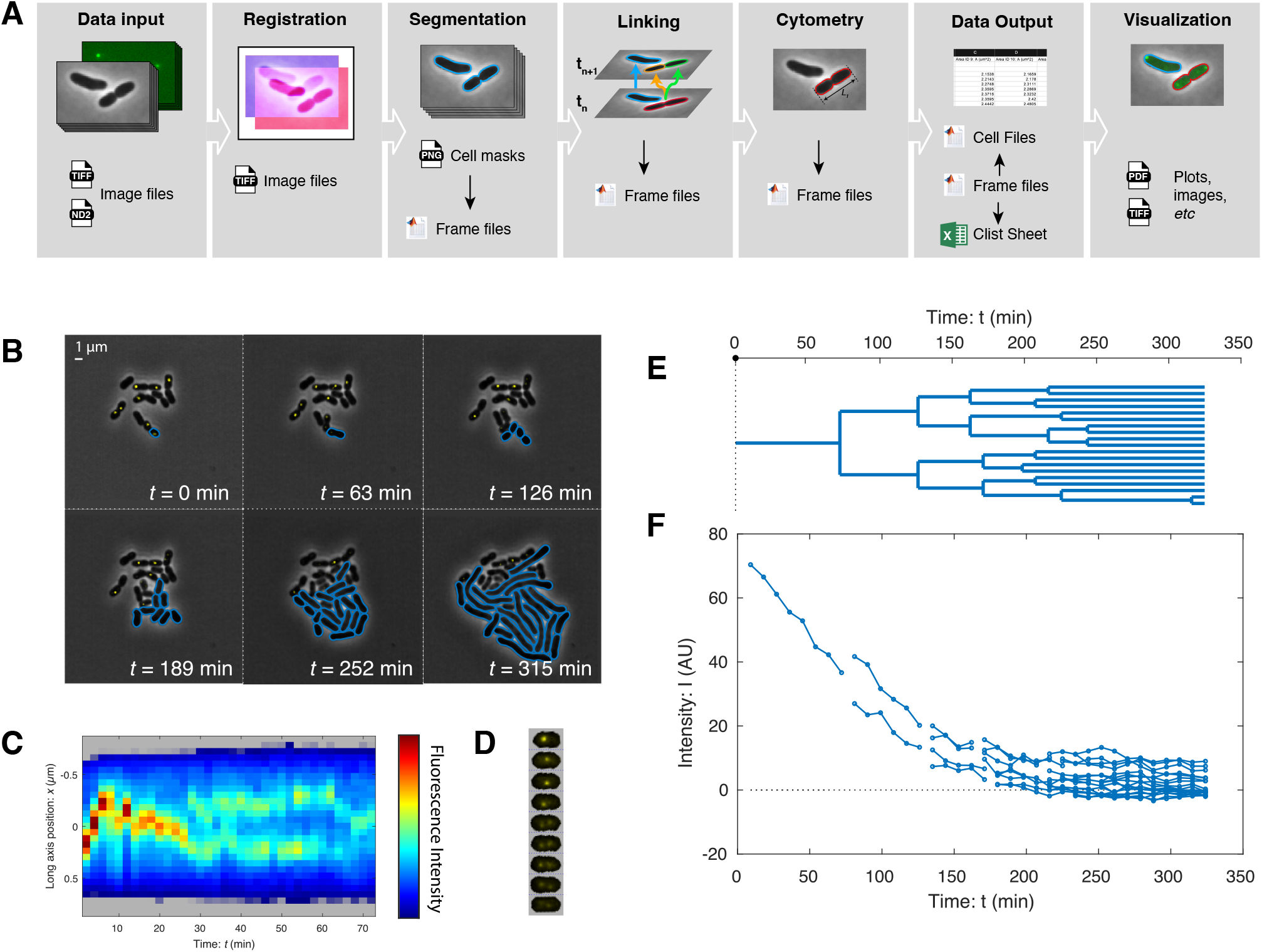
OmniSegger: A microscopy analysis pipeline featuring a wide range of tools for quantitative analysis. **Panel A: OmniSegger pipeline schematic. Data input:** Multi-dimensional image data is loaded from image files. **Registration:** Images are registered to remove stage drift. **Segmentation:** For each position and time-point, the first channel image is segmented to generate the cell masks, which are saved as PNG files. These masks are then incorporated into a frame file, which is a composite data file containing all image information (all channels and cell masks). The cell masks PNG is editable. **Linking:** Cell masks from successive time points are then linked to form cell trajectories, including cell division. The masks and links are corrected for temporal consistency and subsequently saved into the frame files. **Cytometry:** Cell cytometry information for each cell is computed from the image information in the frame files. **Data output:** The output data is sliced into three different output formats: The *frame files* contain all information, including images, grouped by frame (*i*.*e*. all cells per time-point and x-y position). The *cell files* contain all information, including images, grouped by cell (*i*.*e*. all time-points per cell). The *clist file* contains all cytometry information (no image information) grouped per x-y position. **Visualization:** The package also contains numerous visualization tools which use the output data to generate figures, images, and plots. **Panels B-F: OmniSegger visualization gallery**. The pipeline includes numerous tools to visualize cellular growth dynamics. **Panel B: Frame mosaics show proliferation at a multi-cellular scale**. OmniSegger can generate multi-channel composite images and supports the use of vector (rather than raster) cell outlines for improved figures. This panel highlights the ability of OmniSegger to analyze cells with a filamentous phenotype. **Panel C: Kymographs show the intracellular dynamics**. The panel shows the long-axis localization of the replisome (labeled by YPet-DnaN) using false color. **Panel D: Cell towers show intracellular dynamics in 2D**. The panel shows the 2D localization of the replisome (labeled by YPet-DnaN). **Panel E: Lineage trees show cell proliferation from a single progenitor. Panel F: Automated cell cytometry facilitates quantitative analysis**. OmniSegger automatically generates over 100 cellular descriptors, including average fluorescence intensity. In this experiment, the targeted protein YPet-DnaN is depleted by cell-proliferation-induced dilution [6].

#### Image data input

The algorithm is designed to process image data with multiple x-y positions, time points, and fluorescence image channels. The algorithm inputs data from image files following a naming scheme used by the Nikon Elements image export tool; in addition, we include a utility that facilitates the conversion of data into this format.

#### Image registration

Although many modern microscopes use encoded stages, samples are still subject to undesired motion on the micron scale, called *drift* or *jitter*. To minimize the effect of drift, we apply an optional image registration step, on a user-selected channel, to align successive images with sub-pixel resolution using a cross-correlation registration algorithm [7]. We find this step greatly improves both the time-lapse movies generated for visualization, and the robustness of the cell linking analysis.

#### Cell segmentation

The automated detection of the regions in an image corresponding to individual cells is called *instance segmentation* (segmentation) [8]. In the pipeline, the segmentation algorithm works on the first channel of the multi-channel images, and now supports the segmentation of multiple imaging modalities. OmniSegger implements our Omnipose package [3], introducing significant improvements to segmentation performance in comparison to our original release, SuperSegger. The Omnipose algorithm uses a flexible machine-learning-based approach that is well suited to both the problem of segmenting unusual cell morphologies and the segmentation of alternative imaging modalities. The updated algorithm outputs cell masks as PNG files that can be manually edited if segmentation errors arise.

#### Expanded masks

The original SuperSegger package had an important limitation in defining cell boundaries: It required a pixel gap between neighboring cells to define the cell masks; however, when cells grow in micro-colonies, they are typically in contact. For example, for a pixel size of 100 nm and cell area of 3 *μ*m^2^, if we assume boundaries on the cell edges are receded by at least one pixel on each edge, the mask will fail to capture about 10% of the cell, even in the absence of segmentation errors. The original segmentation predicted binary masks for each cell (semantic segmentation); Omnipose predicts uniquely labeled cell masks (instance segmentation). Therefore, masks created in the updated pipeline can be in contact, eliminating this boundary artifact. The improved boundaries defined by the cell masks enable more accurate and consistent measurements of cell properties on the sub-cellular scale (see Fig. 2).

**FIG. 2.**
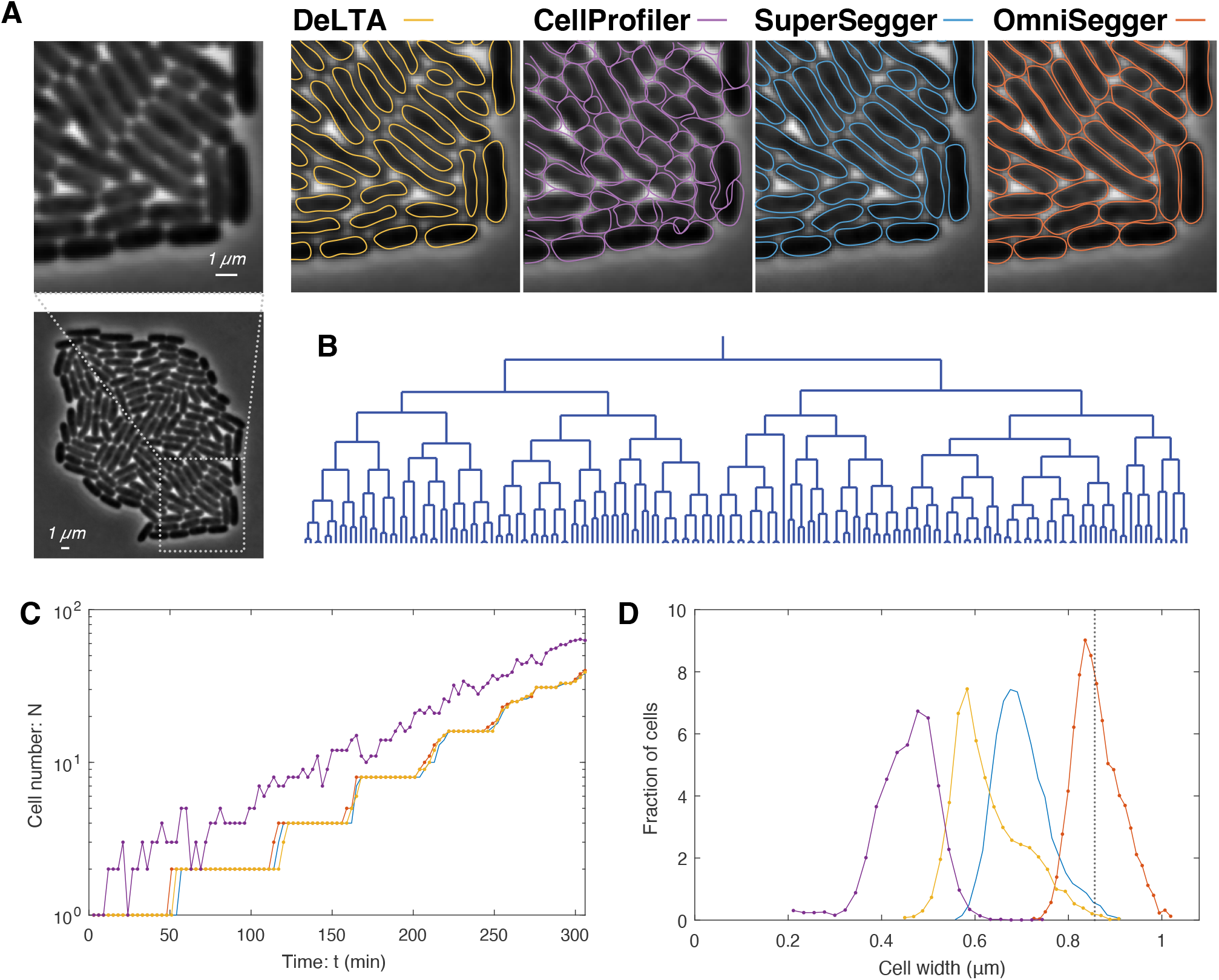
Proliferation challenge. To characterize the performances of the competing packages, we first analyzed performance under the best case scenario: the proliferation of cells, with normal morphology, from a single cell to a monolayered microcolony under ideal imaging conditions. **Panel A: Competing pipelines generate distinct cellular boundaries**. The panel shows a phase contrast image of a microcolony. Competing cell segmentations are shown for a representative magnified region. All the pipelines, except CellProfiler (purple), lead to the same number of cells and are therefore acceptable for colony-scale analysis. The remaining pipelines generate significantly different cell boundaries at a sub-cellular resolution. **Panel B: Lineage tree** as generated by OmniSegger. **Panel C: Quantitation of cell number**. CellProfiler (purple) over-segments the cells to such a great extent that it roughly double counts cells. OmniSegger, SuperSegger, and DeLTA all show comparable performance. **Panel D: Sub-cellular structure**. Panel A visually illustrates the difference in the segmented cellular boundaries. To emphasize the biological significance of these differences, we generated histograms of cellular width measured by each pipeline and compared these to the true average cellular width (dotted line, inferred from cell contact). OmniSegger both generates a measurement with the smallest bias (1%) as well as the narrowest distribution (*σ/μ* = 6%).

#### Linking/tracking

The segmentation algorithm defines cell regions at each time point of a time-lapse experiment; however, to study cellular dynamics, the regions in successive time points must be linked to form cell trajectories. From the labeled mask image, each label assigns a region number to a cell, which is determined on a per-frame basis–region numbers are not persistent throughout the time-lapse. Once the regions have been linked, unique IDs can be assigned to each cell. These cell IDs are distinct from cell region numbers. IDs persist throughout the entire time-lapse experiment.

We provide an alternative version of OmniSegger supporting an updated linking algorithm which is more robust to tracking cells, including those with unusual morphologies. This version of OmniSegger facilitates the use of external cell-tracking packages as well as the hand-correction of linking. The details are discussed in Supplementary Material Sec. B 2.

#### Cytometry

Once the cell IDs have been assigned, OmniSegger performs cell cytometry by computing a user-specified set of cell properties, including cell length, area, birth and division (death) times, mother and daughter IDs, fluorescence intensities, fluorescent-focus position, etc. The current package implements more than 80 default cell characteristics. This cytometry step is the backbone of many analyses, and its accuracy depends sensitively on the precision of preceding steps in the pipeline.

#### Data output

A critical feature of an analysis pipeline for many users is the output of the data in a readable and easily-manipulated format. We provide several data output types that slice time-lapse data in different ways. For instance, we provide data organized by time-point (Frame files), by cell (Cell files), and holistic experimental summary (a clist file), each of which are suited to different analytical tasks. To increase the usability of the pipeline for users not familiar with MATLAB, we now output the clist file in Excel format (in addition to a mat file). This file contains *>* 80 descriptors on a per-cell basis and *>* 20 on a per-cell per-time-point basis, and is easy to expand by the user.

#### Data visualization

To accompany the pipeline, we provide a data-exploration GUI which itself can be used to launch more specialized visualization tools, designed to analyze a wide range of phenomena from localization dynamics in a single cell to comparing population statistics between multiple populations. Each of these tools is provided as MATLAB function to facilitate their adaptation to different applications. One of the new features is the inclusion of improved vector-based cell boundary representations to produce more compelling print figures for publication.

#### Omnipose versus OmniSegger

We recently described the Omnipose segmentation package [3] but we are now releasing OmniSegger. Is OmniSegger better than Omnipose? In short, Omnipose is a part of the OmniSegger pipeline. Like all other parts of the pipeline, it could be used independently; however, for most analyses additional steps are required to convert the Omnipose analyses into interpretable data. OmniSegger performs these additional steps in a package that is designed for bacterial analysis.

#### Additional updated features

In addition to the changes we have described above, we have made numerous additional changes to the OmniSegger pipeline. We provide a detailed list of feature updates in the Supplemental Material Sec. A.

### Measuring performance

When we described the Omnipose segmentation algorithm, we reported performance using the metric of Intersection Over Union (IOU) [3]. Although the IOU metric is widely used when measuring segmentation performance, it combines many different types of errors into a single number. For instance, (i) consistently underestimating the cell width and (ii) erroneously dividing a cell into two regions can both lead to the same reduction in average IOU; however, they affect downstream analysis in very different ways. For instance, erroneous division typically requires hand correction before the pipeline can link the respective regions. We will therefore use two more specific metrics of error in our analysis: We analyze *cell width* as a proxy for the accuracy of sub-cellular scale cell boundaries and define *fatal errors* as over- or under-segmentation errors that require hand-correction before datasets can be temporally linked with accuracy. We will count the cumulative fatal errors during the analysis of the time-lapse dataset as a function of time. This metric has a practical interpretation: this represents the number of corrections that must be made by hand for a quantitative cell proliferation analysis. It is important to point out that these categories are not inclusive of all segmentation errors. For instance, in a time-lapse analysis, if a pipeline detects a cell division a frame early, as long as the regions can be correctly linked, we do not count this early over-segmentation as an error.

### Competing pipelines

We will characterize the new OmniSegger pipeline, and three competing pipelines: DeLTA [9], Ilastik-CellProfiler [10, 11], and SuperSegger [4] in each challenge. The first two pipelines constitute the only complete analysis pipelines that focus on single cell cytometry, require minimal user input, and are actively maintained. Though no longer maintained, we include SuperSegger to demonstrate the improvements of OmniSegger. Although packages such as FAST, Cell-Shape, Oufti, and MicrobeJ exist for time-lapse analysis, these competing packages were not included in the comparison because their segmentation methods are limited or require significant parameter tuning for each dataset [12–15]. The features of the packages are summarized in Tab. I.

### Proliferation challenge

First, we will establish the performance of the pipelines on ideal phase-contrast time-lapse data. This performance will represent an upper bound on the performance.

#### Description of the challenge

We imaged *E. coli* proliferating from a single cell to a microcolony containing more than 100 cells. (See Fig. 2A.) It is important to emphasize that the proliferation challenge is highly non-trivial in two respects: (i) Nearly all algorithms perform well when there is minimal cell contact; however, once the microcolony grows to tens of cells in size, the shade-off phase-contrast artifact leads to reduced cell contrast in the middle of the colony. With this limitation in mind, we were careful to select tightly cropped data where the cells remain tightly focused throughout the time-lapse experiment, and we stop analysis immediately before the microcolony becomes multilayered. To prolong this period for as long as possible, we grew the cells on a 4% agarose pad as previously described [4]. (ii) The area of the colony is expanding exponentially, and therefore the speed of the edges of the colony grow exponentially as well. Tracking cells for time-lapse linking (Fig. 1A) is complicated by this rapid movement of cells at the boundary of the colony at late times. With this limitation in mind, we imaged the micro-colony every 3 minutes to facilitate cell tracking late in the experiment. (The raw images and OmniSegger masks are provided in the Supplementary Material.)

#### Pipeline performance

OmniSegger, SuperSegger, and DeLTA all process the data without fatal errors. (See Fig. 2A.) CellProfiler, which uses a threshold-based segmentation algorithm on Ilastik probability maps, has such poor performance as to make automated analysis of this dataset intractable: The pipeline overestimates the number of cells by about a multiple of two throughout the time-lapse (Fig. 2C).

Except for CellProfiler, the other three pipelines process the data without fatal errors, however, there are significant differences in their performance at a sub-cellular scale. Fig. 2D shows a quantitation of cell width measured by each pipeline compared with the true cell width as inferred from cell contact in the microcolony. (See Supplemental Material Sec. C.) We summarize this performance in Tab. II. OmniSegger shows the smallest bias and cell-to-cell variation in cell width. The differences between the package performances are significant, and we therefore conclude that OmniSegger is the only package suitable for analyses where sub-cellular scale resolution is important.

### Morphology challenge

The segmentation of bacterial cells with unusual morphologies was a critical weakness of our original SuperSegger pipeline and a central motivation for the current work. We have encountered many experimental scenarios in our own projects where the pipeline failed, including our attempt to characterize essential-gene deletions in the model bacterium *Acinetobacter baylyi* [16] where the performance of the SuperSegger pipeline was so poor that we were forced to abandon the original aim of our experiments until we developed the OmniSegger package [6].

#### Description of the challenge

To measure the performance of competing pipelines on unusual cell morphologies, we analyze time-lapse images of cells forming filaments which traditional pipelines typically oversegment. We therefore collected time-lapse data of *E. coli* cells treated with a sub-Minimum Inhibitory Concentration (sub-MIC) of 10μM hydroxyurea [17, 18]. Hydroxyurea inhibits DNA synthesis, and results in a phenotype of cell filamentation when below the MIC. See Fig. 3A. (The raw images and OmniSegger masks are provided in the Supplementary Material.)

**FIG. 3.**
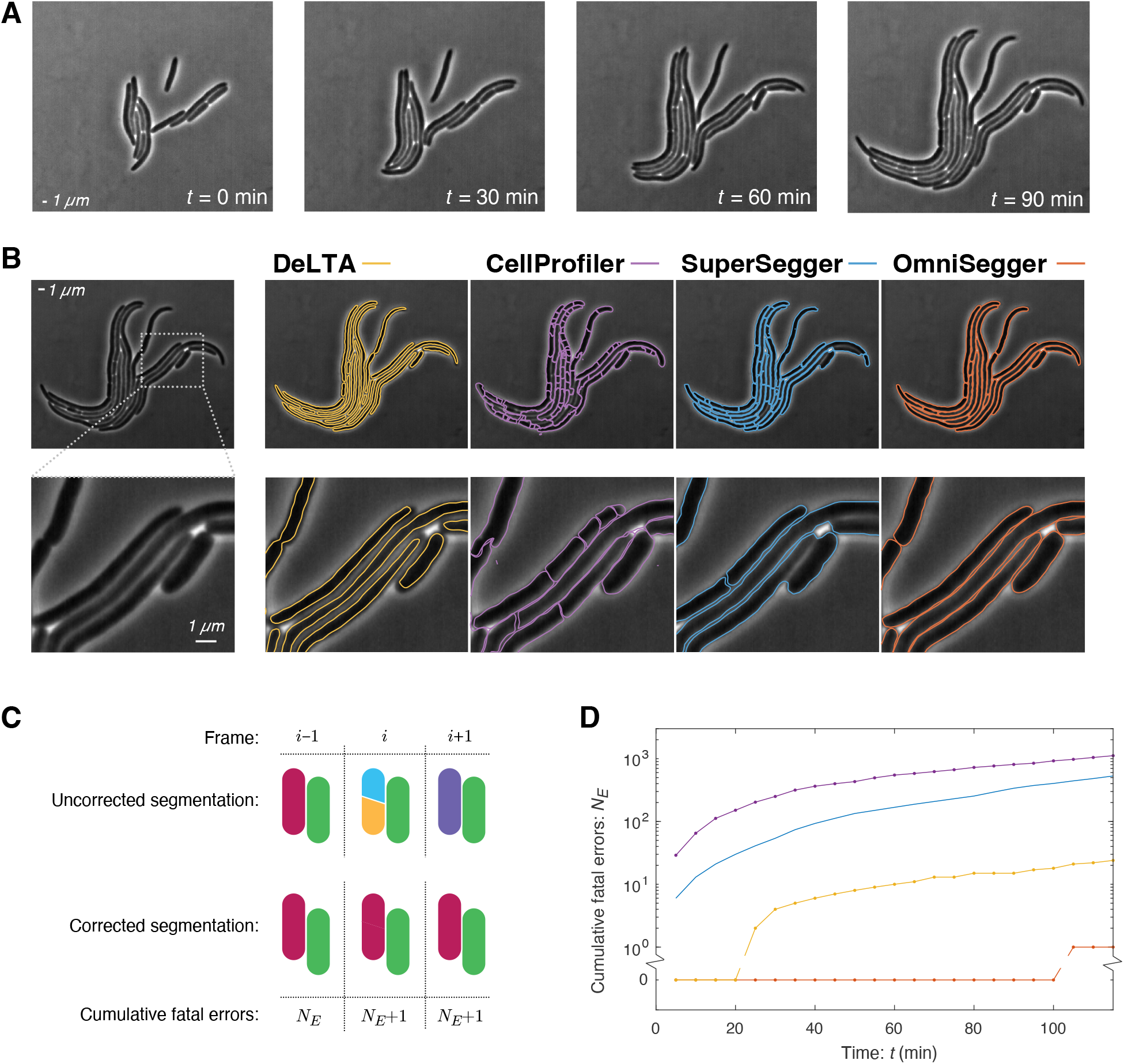
The performance of competing pipelines on unusual cell morphologies. **Panel A: Visualization of proliferation.** Frames from a phase-contrast time-lapse of a growing wild-type *E. coli* colony treated with a sub-MIC of hydroxyurea. Hydrox-yurea inhibits DNA synthesis and results in a phenotype of cell filamentation. **Panel B: Competing pipelines generate distinct cellular boundaries**. Pipeline performance varies greatly for cells with unusual morphologies, not only at sub-cellular resolution, but at a colony scale. For this challenge, we focus on *fatal errors* defined as those that affect the cell number and prevent temporal linking without hand correction. **Panel C: Cumulative fatal error defined**. We measure performance as cumulative number of fatal errors. The red cell is over segmented in the *i*th frame, generating two new cells (yellow and cyan) before fusing back into the original cell (purple). The corrected segmentation is shown below. This segmentation error in frame *i* increases the cumulative error *N* by 1. Note that neither erroneously narrow cell boundaries nor a late (or early) call of a cell division event constitutes a fatal error. **Panel D: Performance of competing packages measured by cumulative fatal errors**. The OmniSegger analysis is error free for 100 min of imaging (20 frames). DeLTA also results in a tractable analysis, although the analysis requires the correction of over 20 cells in a single microcolony. The performance of the SuperSegger and CellProfiler pipelines are so poor for cells of unusual morphology as to make these analyses intractable.

#### Pipeline performance

The analysis of unusual cellular morphologies led to a wide range of pipeline performance. The segmentation of a representative frame is shown in Fig. 3B. A schematic definition of fatal errors is shown in Fig. 3C and a plot of cumulative fatal errors as a function of time in shown in Fig. 3D.

CellProfiler had the worst performance. The package leads to both chronic over- and under-segmentation and generates a completely intractable number of fatal errors. Although SuperSegger performed well on normal cell morphologies, it had extremely poor performance for filaments and generated an intractable number of fatal errors. DeLTA again fails to accurately capture cell width; however, it performs well with respect to fatal errors. For the whole dataset, only 23 fatal errors would need to be fixed, which we would define as tractable. The performance of OmniSegger is, by both performance metrics, the best. The cell boundaries are accurate at a sub-cellular scale (Fig. 3B) and there is only one fatal error in the entire time-lapse which occurs very late in the experiment (Fig. 3D). We therefore conclude that OmniSegger has the best performance in the context of unusual cell morphologies.

### Modality challenge

The second focus of OmniSegger development was to add support for the segmentation of more imaging modalities. Although we prefer to use the phase-contrast modality in most experiments, some experimental scenarios demand the use of other imaging modalities. For instance, phase-contrast objectives contain a neutral-density ring that decreases the overall brightness of the objective in fluorescence applications, making it a sub-optimal modality for experiments that demand single-molecule sensitivity. We therefore worked to create a pipeline with robust segmentation performance using any of a range of canonical imaging modalities. Due to our own laboratory priorities, we introduced support for the following modalities: Phase-contrast, Brightfield, cytoplasmic fluorescence, and membrane fluorescence.

*Description of the challenge*. For this challenge, we collected data using four imaging modalities (Phase-Contrast, Brightfield, cytoplasmic and membrane fluorescence). In each case, we collected time-lapse data of *E. coli* cells proliferating over multiple generations. *For the phase-contrast modality*, we used the dataset as described in the Proliferation challenge. *For the brightfield modality*, we imaged cells using a z-stack, but focused on in-focus and under- and over-focused (± 0.5 μm) images. We also experimented with two growth conditions: rich (LB) and minimal (M9). We did find significant differences in brightfield images and segmentation performance between these conditions and focused on the minimal media data for which the segmentation performance was highest. *For the fluorescence modality*, the performance is clearly dependent on the brightness of the fluorescent label. We assumed that most users would desire to use plasmid-based fluorescent fusions for this purpose. We therefore selected two representative plasmid-based IPTG inducible fusions from the ASKA collection: One with diffuse cytoplasmic localization and the other with membrane localization [5]. (The raw images and OmniSegger masks are provided in the Supplementary Material.)

#### Pipeline performance

While segmentation algorithms and pre-trained models exist to segment various imaging modalities, among the currently existing quantitative time-lapse analysis pipelines, the majority only support segmentation for phase-contrast images. In addition, alternative pipelines, which are able to segment brightfield or fluorescence images, require significant time investment to tune user-defined training or parameters (see Table I). OmniSegger is the only pipeline which can handle various imaging modalities out-of-the-box. As shown in Fig. 4, the OmniSegger pipeline can handle all five modalities. Since other packages either do not support these alternative modalities (DeLTA, SuperSegger) or have extremely poor performance (CellProfiler), we show only the OmniSegger results. In order to evaluate the relative performance with each modality, we again quantified the cumulative number of fatal errors for cells propagating from single cells to form microcolonies (Fig. 5).

**TABLE I.** Analysis pipelines & features. A comparison of features and functions for cellular image analysis software packages.

| Pipeline | Tuning | Segmentation | Tracking | Data | Output | Coding | OS |
| --- | --- | --- | --- | --- | --- | --- | --- |
|  | free | algorithm | algorithm | visualization | format | Language | support |
| OmniSegger | ✓ | Deep learning (Omnipose) | Traditional | ✓ | mat, xls | MATLAB & Python | Linux, Windows, MacOS |
| SuperSegger [4] | ✓ | ML-informed Threshold | Traditional | ✓ | mat | MATLAB | Linux, Windows, MacOS |
| DeLTA [9] | ✓ | Deep learning | Deep learning | ✗ | nc | Python | Linux, Windows |
| Ilastik-CellProfiler [10, 11] | ✗ | ML-informed Threshold or Watershed | Traditional | ✗ | xls | Standalone | Linux, Windows, MacOS |
| FAST [12] | ✗ | Threshold | Unsupervised learning | ✓ | mat | MATLAB, Standalone | Linux, Windows, MacOS |
| CellShape [13] | ✗ | Threshold | N/A | ✓ | N/A | Python | Linux, Windows, MacOS |
| Oufti [14] | ✗ | Threshold | Traditional | ✓ | mat, out, csv | MATLAB | Linux, Windows |
| MicrobeJ [15] | ✓ | Threshold | Traditional | ✓ | res, csv | Java (ImageJ) | Linux, Windows, MacOS |

**TABLE II.** Precision in determining cell width. We use cell width as a proxy for sub-cellular scale segmentation resolution. Comparison of bias and standard deviation in determining cell width are shown for each pipeline. Only OmniSegger generates cell boundaries precise enough for applications which depend on precise cell boundary position. OmniSegger also measures cell width with the smallest cell-to-cell variation.

| Pipeline: | Bias: $\Delta\mu$ | | Std: $\sigma$ | |
| --- | --- | --- | --- | --- |
|  | (%) | (nm) | (%) | (nm) |
| OmniSegger | 0.6% | 5 | 6% | 50 |
| SuperSegger | -18% | -160 | 7% | 60 |
| DeLTA | -27% | -230 | 9% | 80 |
| Ilastik-CellProfiler | -47% | -400 | 8% | 70 |

**FIG. 4.**
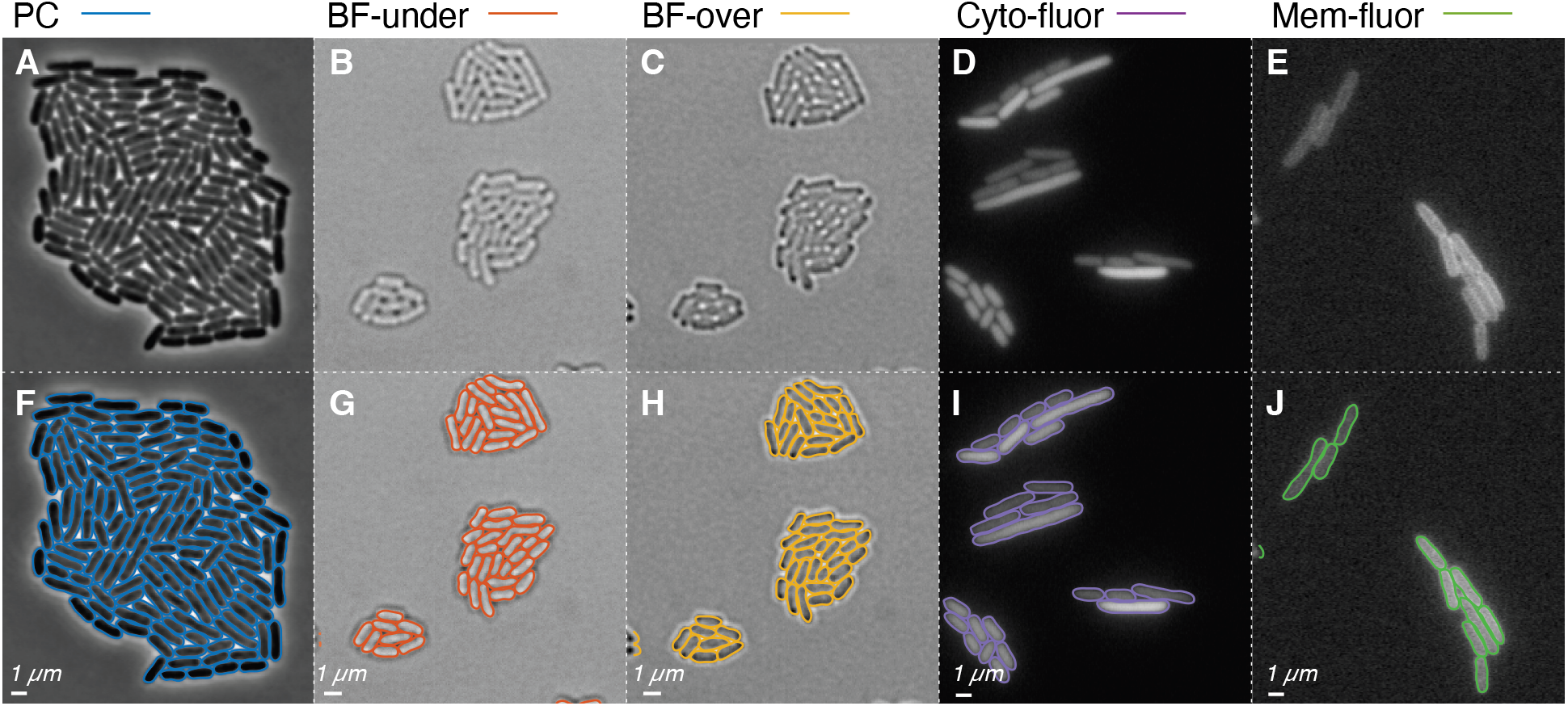
OmniSegger robustly segments multiple imaging modalities. **Panel A & F: Phase-contrast** (PC) imaging results in high-contrast images with well-defined cell boundaries. We find this modality generates the highest performance segmentation. **Panels B, C, G, & H: Under-focused brightfield** (BF-under) and **Over-focused brightfield** (BF-over) both generate low-contrast, noisy images. After training on this modality, we found the performance of OmniSegger to be high enough for many applications, although PC is still superior for high cell density. We also analyzed in-focus brightfield images; however, performance was not high enough to recommend this modality. **Panel D & I: Cytoplasmic fluorescence** labeling (Cyto-fluor) results in high-contrast but noisy images with bright cell interiors. We found that the Cyto-fluor modality could lead to excellent performance; however, this modality is subject to photobleaching and phototoxicity which limits the signal-to-noise ratio in some applications. **Panel E & J: Membrane fluorescence** labeling (Mem-fluor) generates high-contrast but noisy images with bright cell boundaries with nearly uniform cytoplasmic labeling from the out-of-focal-plane membrane. We found this modality could lead to high-performance segmentation; however, it was particularly sensitive to noise, especially when compared with Cyto-fluor.

**FIG. 5.**
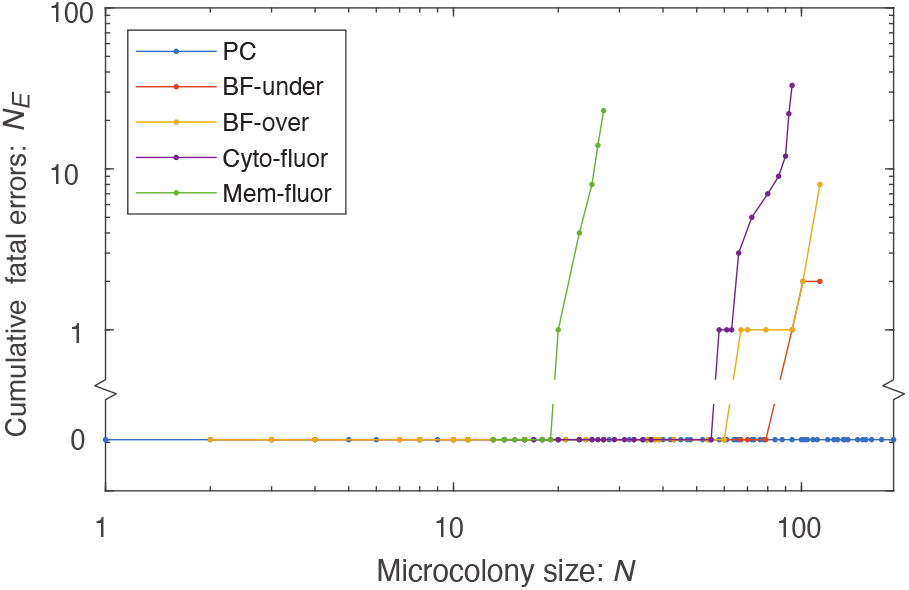
OmniSegger performance for a range of image modalities. In each case, cell proliferation was visualized on agarose pads, starting from single isolated progenitor cells growing to form microcolonies. Differences in cell growth arose from the use of distinct growth conditions and strains, therefore we quantified the fatal error number versus colony size (in cells). Phase-contrast data were analyzed without fatal errors. Over- and under-focused brightfield imaging led to high-performance segmentation as well, followed by cytoplasmic fluorescence labeling. Membrane fluorescence labeling led to the worst performance of all modalities; however, we emphasize that the performance is sufficient for many applications.

For the challenge, we chose datasets that were tractable and led to high segmentation performance. In this context, all modality data were processed without fatal errors until the microcolony reached 20 cells, where Membrane-fluorescence began to fail. Cytoplasmic-fluorescence demonstrated superior performance compared to membrane-fluorescence. Furthermore, performance on both under-focused and over-focused bright-field surpassed that of cytoplasmic fluorescence. Nevertheless, phase-contrast had the fewest fatal errors among all the aforementioned modalities. It is important to emphasize that for many applications, the performance of all modalities is sufficient to extract quantifiable data. However, we note that this challenge underestimates the advantages of phase-contrast over the competing modalities.

#### Brightfield performance

Although the performance of OmniSegger on the brightfield images is high in the challenge, it is important to discuss a number of caveats about these results. (i) Notably, we were unable to generate a high-performance algorithm for in-focus data in spite of extensive training. We concluded that the contrast was too low relative to the competing mechanism of contrast generation. Using the brightfield modality therefore requires a focal plane shift relative to fluorescent images, which, while tractable, is not ideal. (ii) We observed significant differences in brightfield segmentation performance depending on the growth conditions. We observed high performance on minimal media but lower performance on LB. This drop in performance was consistent with our intuition of the modality image quality when comparing the images by eye: The minimal media data was clearly more interpretable. In comparison, we find the image quality and segmentation performance for phase-contrast is much less dependent on growth conditions. We therefore conclude that the brightfield modality segmentation may have excellent performance in many contexts; however, it may be somewhat inconsistent and dependent on the dataset.

#### Fluorescence performance

There is good news for fluorescence based modalities as well: OmniSegger can segment data with very high performance. However, like the brightfield modality, there are important caveats. Clearly, both fluorescent modalities depend upon label brightness and fluorescent labels are typically subject to bleaching, which can significantly attenuate their brightness. As the intensity decreases, we found that cytoplasmic labels tend to lead to more robust performance than membrane labels. Our intuition for this difference in performance is that the contrast for the membrane label tends to be very flat across the cell, whereas the cytoplasmic label gives rise to a significant gradient towards the cell boundaries, which is more robust to noise. (See Fig. 6.) In conclusion, the fluorescence modality can lead to high performance segmentation, and the cytoplasmic label is preferable.

**FIG. 6.**
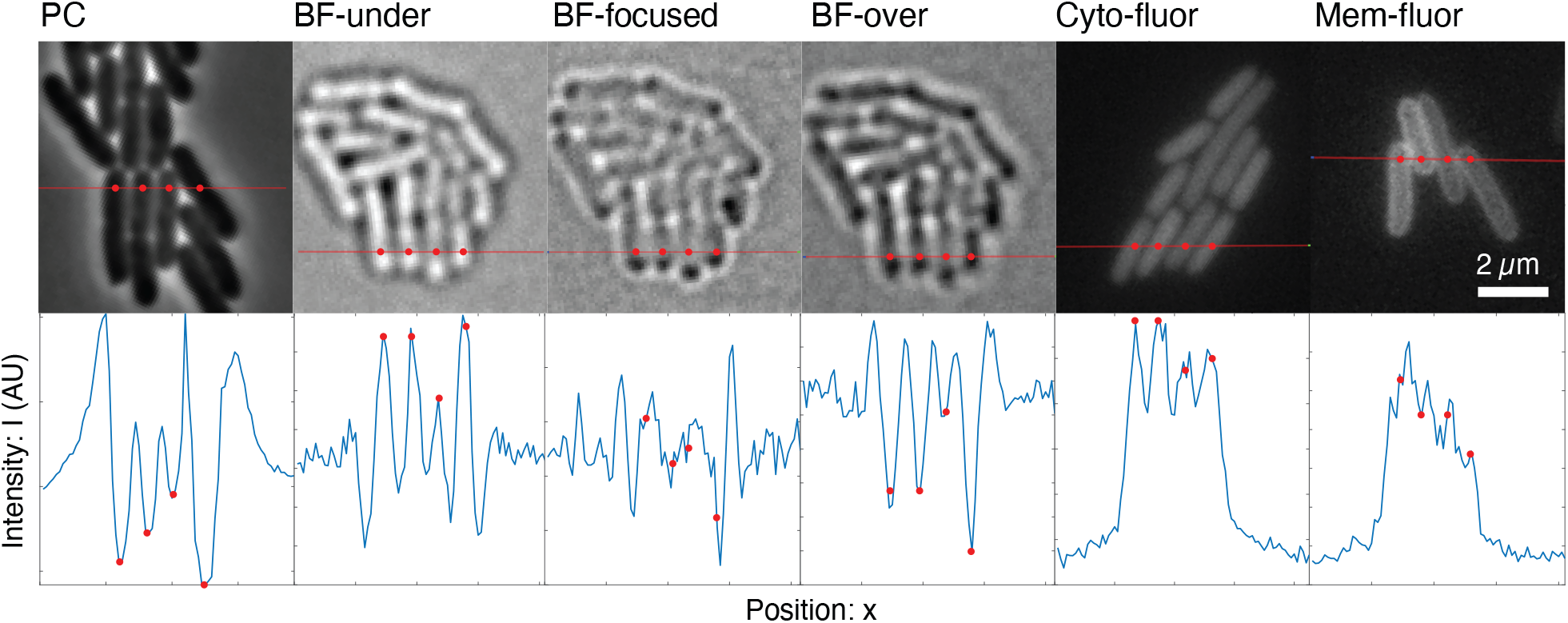
Contrast in a range of modalities. **Top row: Images for a range of modalities.** For each modality image, we chose a line that transects multiple cells. **Bottom row: Line scan**. The line-scan plot shows the intensity profile across the red line above. In addition, we annotated a number of points corresponding to cell centers (red points). In the modalities where the segmentation performance in highest (PC, BF-under, BF-over, Cyto-fluor), there are clear morphological features in the line-scan plot corresponding to the cell; however, in the lower performing modalities (BF-focused and Mem-fluor), the morphological features corresponding to cells are less distinct, making these modalities susceptible to contamination by noise.

## III. DISCUSSION

### Morphological robustness enables new approaches

The development of OmniSegger enables the quantification of a variety of novel experiments involving mutants, antibiotic-treated cells, and multi-species interactions, at the single cell level. Many of these experiments involve observing a wide range of cell morphologies. Due to the challenge of accurate and robust cell segmentation, these analyses were previously not possible. The development of Omnipose enabled the segmentation step of the image analysis pipeline, and its implementation in OmniSegger introduces a powerful, highly automated tool for the biological imaging community.

### Expanded modality support offers greater flexibility

Cell biology experiments routinely require a mix of approaches, including the use of different imaging modalities. The robust performance and analysis capabilities of OmniSegger, irrespective of the modality used, are greatly superior to other packages. Even if alternative packages could match the performance for a single modality, the generalist support offered by OmniSegger enables a uniformity in approach to multiple experiments and the advantage of avoiding package-dependent biases into an analysis.

### Understanding modality-dependent performance

Can the observed differences in segmentation performance between modalities be rationalized based on image contrast? In Fig. 6, we compare the intensity profiles from a line scan in each imaging modality. At an intuitive level, we observe that each of the modalities that is segmented with high performance (phase-contrast, under- and over-focused brightfield, and cytoplasmic fluorescence) all have consistent morphological features corresponding to each cell. In comparison, the other poor-segmentation-performance modalities (in-focus brightfield and membrane fluorescence) have less consistent morphological features distinguishing neighboring cells. We therefore conclude that the relative algorithmic performance is consistent with what we would intuitively predict based on the ability of the image modality to generate unambiguous image contrast at cell boundaries.

**TABLE III.**
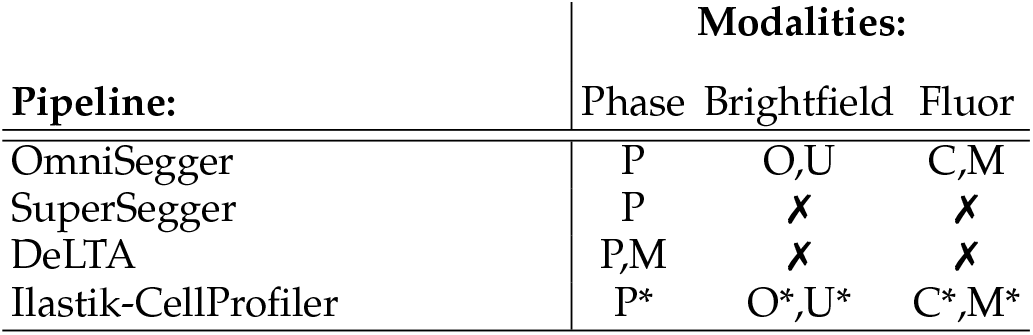
Pipeline modality support. For phase-contrast, **P** denotes agarose pads and **M** denotes mother machine. For brightfield, **O** denotes over-focused and **U** denotes underfocused. For fluorescence, **C** denotes cytoplasmic and **M** denotes membrane labels. * denotes tuning required.

### Which modality?

Given the flexibility to choose from multiple imaging modalities for an experiment, which modality enables the most accurate analysis? Our initial preference, based on our past work, was phase contrast imaging. Is this preference still supported in light of OmniSegger performance? Phase-contrast is, without doubt, still the best modality for determining cell boundaries for bacterial cells. In our own experience, our analysis underestimates the relative advantages of this approach. We selected datasets, growth conditions, and exposures to place the segmentation performance of each modality in the best light possible. However, in our experience, phase-contrast imaging leads to robust and reliable results with suboptimal images and it is therefore our go-to choice for imaging bacteria for quantitative analysis.

### Performance of competing packages

We have provided extensive evidence of the performance advantages of OmniSegger over competing packages in three challenges. The machine-learning approach used in OmniSegger gives it a particular performance advantage over threshold-based pipelines [10, 13–15, 19, 20] that are particularly sensitive to imaging conditions and morphology. Although some competing pipelines can theoretically generate comparable single-cell cytometry, including the generation of cell lineages and mean fluorescence levels [9, 12, 21], their poor segmentation performance limits their applicability. In other cases, packages provide high-performance segmentation but lack a complementary pipeline essential for experimental analysis [2, 11, 22]. A summary of features for OmniSegger and alternative packages are shown in Table I and more extensively in Table S1. In summary, OmniSegger provides (i) high performance segmentation and (ii) the analysis pipeline required to convert these regions into a quantitative analysis with minimal user input or coding experience required.

## Data availability

*Brightfield model*. The Omnipose brightfield model is available on Zenodo […] under the CC-BY-NC 4.0 license. *Brightfield datasets*. Bacterial brightfield and fluorescence image sets of *Escherichia coli* and *Burkholderia thailandensis* were generated in this study. Additional *E. coli* and *Staphylococcus aureus* brightfield images were sourced from a selection of the DeepBacs datasets [23–25] which is available under the CC-BY 4.0 license. The brightfield images, fluorescence images, and ground truth masks used to train the brightfield Omnipose model are available on Zenodo […] under the CC-BY-NC 4.0 license. *Analysis datasets*. Image and OmniSegger mask files of the data presented in this paper are available on Zenodo […] under the CC-BY-NC 4.0 license.

## Code availability

*OmniSegger package*. The code for OmniSegger is available on GitHub: https://github.com/tlo-bot/omnisegger/.

## Acknowledgments

The authors would like to thank S. Yang, B. Traxler, S. van Teeffelen, and J. Mäkelä. H.J.C., T.W.L., and P.A.W. were supported by NIH grant R01-GM128191 and NSF grant GR046955. K.J.C. was supported by the Molecular Biophysics Training Program (NIH grant T32GM008268).

## Author contributions

T.W.L., H.J.C., K.J.C., and P.A.W. conceived the research. T.W.L., H.J.C., K.J.C., and P.A.W. performed the experiments. T.W.L., H.J.C., and P.A.W. performed the analysis. T.W.L., H.J.C., K.J.C, and P.A.W. wrote the paper.

## Supplementary Materials

Supplementary text, Materials and Methods

Table S1.

Data S1-S7.

## Supplementary Material

## Appendix A: Feature updates: From SuperSegger to OmniSegger

OmniSegger skips the original SuperSegger cell segmentation algorithm, and runs the Omnipose package. The SuperSegger mask variables are replaced by the masks output by Omnipose. Omnipose segmentation is far more robust to diverse cell morphologies in addition to improving performance on a subcellular scale. As a result, Omnipose provides the foundation for time-lapse analysis by OmniSegger. In addition, the data visualization features have also been updated to accommodate improved segmentation results.

### 1. Omnipose segmentation

Omnipose is a deep neural network (DNN)-based algorithm for cell segmentation [3]. It is originally based off of the Cellpose algorithm [26]; however, it makes several changes which significantly improve performance.

The use of the distance field, suppressed Euler integration, and training a phase-contrast model on a diverse dataset allows Omnipose to be a leading segmentation tool. Omnipose is much more robust to various imaging conditions and cell morphologies than other cell segmentation algorithms [3].

### 2. Improvements in data visualization

We introduce new data visualization ideas and improvements to generalize for diverse cell morphologies: 1) cell outlines, 2) medoid for cell ID display, and 3) figure making tools.

#### 1. Cell outlines

The displayed cell outlines were previously calculated by dilating the masks. The outlines are now calculated using a more robust cell perimeter function directly based on the mask (`getperim.m`).

#### 2. Medoid of cell skeleton for cell ID display

Many cell analysis software packages, including the original SuperSegger, determine the centroids of cell masks as an approximation for the cell ‘center’ and use the centroid in order to perform measurements. A centroid is a point calculated from the mean position of all data points, *i*.*e*., the mean pixel position of the cell mask. For a given number of *N* pixels with the coordinate of pixel *i* given by *x*_*i*_, *y*_*i*_, the centroid 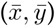 is simply calculated by:

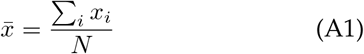

and

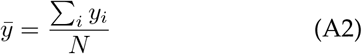

In MATLAB, this is calculated using the `regionprops` function.

The centroid approximation for the cell ‘center’ fails for more diverse morphologies, for example with filamented cells that contain curvature, where the centroid would fall outside the mask. Rather than calculating the centroid, OmniSegger determines the *medoid*. The medoid is the position of a pixel contained in the cell mask which has the minimum sum of distances from every other pixel in the mask:

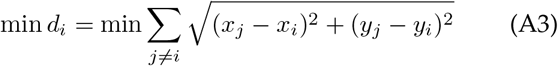

Furthermore, restricting the medoid to the skeleton results in an even more intuitive result for the cell ‘center’ (see `find medoid.m`). Using the medoid of the skeleton as the cell ‘center’ displays the cell IDs much more intuitively than the centroid, applicable to both rod-shaped and other morphologies.

#### 3. Figure making tools

Generating publication-quality figures is important for disseminating results of scientific research. OmniSegger includes new functions for generating figures: `getFamily`, which determines a cell lineage from a progenitor cell, `drawCellSpline`, which draws vectorized outlines of cell boundaries on a cell-by-cell basis, and `makeMosaic`, which generates a mosaic image of selected frames from a time-lapse, with the ability to display fluorescence and cell outline overlays.

### 3. Modified cell length measurement

In SuperSegger, the cell length is calculated using MATLAB’s `regionprops` function. `regionprops` fits a cell as an ellipse, and we expect this approximation to be poor for morphologies with curvature. Therefore, we replaced the cell length measurement with a rod length in `rodGeom`.

The area *A* of a cross-section of a 3D rod is approximated by a rectangle of length *L* and width 2*R* with two half-circle end caps of radius *R*:

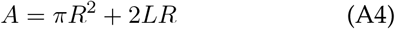

We can define the length of this rod as the length of the rectangle plus with the two end caps:

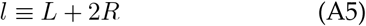

We can also find the integral of the distance field, *B* = *B*_caps_ + *B*_rect_:

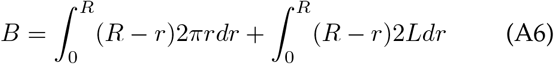

and evaluate to find:

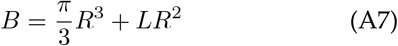

As we want to solve for the rod radius *R* and length *l*, we substitute for *L* in Eqs. A4 and A7 using Eq. A5 to find:

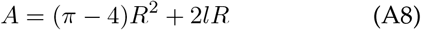

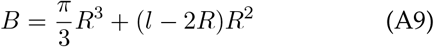

We first solve for *R*. We combine the equations for *A* and *B* to cancel out the terms with *l* resulting in a cubic equation, which we rearrange to the form of a depressed cubic equation:

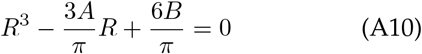

We define two constants *p* and *q* to solve the depressed cubic equation:

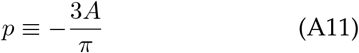

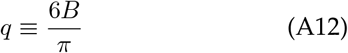

We then calculate a quantity related to the discriminant:

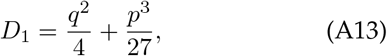

and using Cardano’s formula, calculate the three possible roots for *R*:

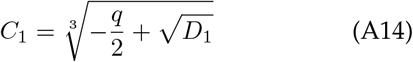

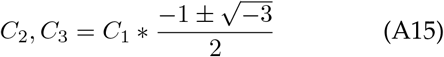

The resulting values are then filtered for only real, positive roots, and the smallest root is chosen as the *rod radius, r*_rod_. The *rod length, l*_rod_, is then calculated by plugging in the rod radius into Eq. A4:

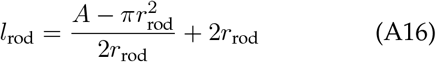

In MATLAB, we calculate *B* using the discrete sum (rather than the integral) of the distance field of the mask. To account for this discrepancy, we subtract an offset term found from simulating pre-defined masks and calculating their rod radius and length:

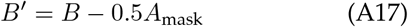

*B*^*′*^ is used when calculating *q* in Eq. A12.

### 4. Improvements in accessibility

#### a. The clist as an Excel spreadsheet

To improve the accessibility and ease of analysis for researchers, we include a new function `clist2xls.m` which allows users to save the clist as an Excel sheet, in addition to its default `.mat` format.

#### b. ND2 to TIFF conversion

We include a Python script `nd2totiff.py` which utilizes the `aicsimageio` package to save ND2 files as individual TIFF files with the OmniSegger naming convention. The script can accommodate multiple channels, XY positions, time-points, and z-planes; in addition, metadata is saved as a TXT file.

## Appendix B: Limitations of OmniSegger

While OmniSegger offers significant improvements to the time-lapse analysis pipeline, in practice, various issues remain. The most pressing issues for OmniSegger at the moment are i) segmentation errors and inconsistent calling of divisions possibly causing cell linking errors, and ii) an open question about the capabilities of cell segmentation-analysis software packages.

### 1. Discussion: Challenges for linking algorithms

The majority of cell tracking algorithms–including OmniSegger–take masks as inputs, then define frame-to-frame linking costs for each cell mask based on properties such as mask overlap, and minimize the costs to determine links. Determining links and segmentation simultaneously is often computationally intensive, as the cost matrix presents an exponentially growing combinatorics problem; if not determined simultaneously, cell tracking accuracy is totally dependent on cell segmentation results. Thus, if the preceding cell segmentation contains errors, the tracking and lineage determination will be disrupted.

Precisely determining the exact time of division for a cell in an experiment is impossible–time-lapses observe the instantaneous event in discrete time steps. Furthermore, the time of division is often ambiguous when only observed by phase-contrast imaging; this was the primary reason for training the Omnipose bacteria phasecontrast model based on underlying membrane or cytosol fluorescence signal when possible. Due to the ambiguous nature of the phase-contrast image, the algorithm will have a ‘flickering’ effect; for example, the cell may be determined to be divided into two cells in frame *t*_0_, but one cell in the subsequent frame *t*_0_ + 1, and back to two cells in frame *t*_0_ + 2. This flickering effect presents an issue for 2D segmentation algorithms, especially when the time-lapses are taken at high frame rate. At the moment, existing 2D segmentation models only perform segmentation by considering individual frames, while the model determination about cell division must persist across multiple frames.

Omnipose introduces the idea of a ‘spacetime’ model, where the segmentation algorithm takes temporal information into account, effectively becoming a 3D model (2D+T). Let’s consider the kymograph for a cell dividing into two daughter cells: the mother cell grows until the cell wall septates roughly near the center. Conceptually, the kymograph appears like pants; as time increases, the mother is the waist of the pants, which splits into two daughter pants legs. The split is persistent in time. Assuming a frame rate high enough that cells overlap from frame to frame, the 3D spacetime segmentation also presents a solution for cell linking, as each 3D lineage volume contains the time of division and the mother-daughter information. While the 3D spacetime Omnipose model (`bact phase spacetime`) is promising, it is lacking the large ground-truth, annotated training dataset as used for the 2D segmentation model and as a result is much less robust.

Though highly robust, Omnipose’s `bact phase omni` used with OmniSegger can have segmentation errors and in addition, its 2D model does not fix the persistent cell division issue.

A unique feature of SuperSegger/OmniSegger is the inclusion of error resolution functions which can edit the underlying masks to solve segmentation and linking errors, though it is also not fully robust.

### 2. Results: Bactrack–An alternative linking algorithm

As previously mentioned, there are various limitations to the original tracking algorithm. We have implemented a form of the *Bactrack* package into OmniSegger, which is offered as an alternative option for cell tracking, specializing in linking for diverse morphologies.

Motivated by observations where the original linking algorithm performed poorly for filamentous cells, Sherry Yang developed the Bactrack package [27] in Python, which provided much improved tracking performance. Bactrack is a cell tracking tool which uses hierarchical segmentation and mixed-integer programming optimization. The hierarchical segmentation approach of Bactrack was inspired by ultrack [28]; Omnipose flow field inputs are used to create hierarchical segmentations from low to high resolution.

Given a graph of possible segmentations, a matrix of linking costs, and the constraint that cells only divide, Bactrack optimizes the final graph of cell segmentations. Bactrack outputs *both* masks in Omnipose labeled-mask format, and linking results as a Pandas dataframe.

Bactrack allows the option to use one of three different MIP solvers: HiGHS [29], and CBC [30] and Gurobi [31] through Python-MIP [32], though Gurobi and HiGHS are the fastest [33]. OmniSegger only implements HiGHS and Gurobi.

While in theory, the masks that result from the optimized hierarchical segmentation should be more accurate than Omnipose segmentation, in practice, the resulting Bactrack masks were observed to be less biologically accurate; the Bactrack masks often called division too early or too late compared to Omnipose masks when tested on the same dataset. Therefore, the OmniSegger implementation inputs Omnipose masks into Bactrack to generate only linking results. Furthermore, the error resolution code in OmniSegger relies on checking linking results. If the linking results are now determined by Bactrack, error resolution by OmniSegger is not possible and thus the version of OmniSegger with Bactrack implementation does not correct the underlying masks. This can be most dramatically demonstrated by a high time-resolution timelapse, which can suffer from inconsistent determination of division from frame-to-frame.

The linking results contain the mappings between a ‘source’ (*t*) frame and a ‘target’ (*t* + 1) frame. The format of results is as follows: the first column lists the source frame starting at frame 0. The second column lists the label IDs in the source frame that have mappings to the target frame. The third column contains the label IDs in the target frame that are mapped from the source frame. In addition, the fourth and fifth columns are source frame cell areas and target frame cell areas, which are used for tracking error calculations. Note that if a cell in a source frame does not have a mapping in the target frame (*i*.*e*., a stray cell which appears in the source frame but disappears in the target frame), there will be no entry for that cell ID in the Bactrack linking results; OmniSegger fills these links in `fillBactrackLinks.m`.

The linking results are then converted from Pandas dataframe into comma-separated values (csv). The csv file is read into MATLAB during the linking stage in OmniSegger and then converted into SuperSegger linking format with forward and reverse mappings. In case of errors, the csv can be manually edited before further linking by OmniSegger.

Bactrack is implemented with OmniSegger through the following GitHub branches: Bactrack branch super-SeggerDev, OmniSegger branch bactrackdev. This version of OmniSegger is recommended when the main branch of OmniSegger fails to track unusual morphologies for timelapses at medium to low frame rates.

### 3. Results: Modular segmentation

Omnipose is not fully implemented into OmniSegger. Instead, its output masks is the input to the segmentation step of OmniSegger. In fact, any other segmentation algorithm can be used, as long as the input masks are in `png` format. The modularity of OmniSegger allows it to be compatible with future advances in cell segmentation. Similarly, the *bactrackdev* branch of OmniSegger introduces modularity in linking, as long as the input `csv` is in Bactrack format.

### 4. Discussion: Generalization of cell cytometry software

While the introduction of modular segmentation and linking steps allows OmniSegger to be much more robust to analyzing time-lapses of diverse cell morphologies, the analysis software is still biased towards rodshaped cells. For example, quantities such as long axis length or cell pole age are calculated. However, consider cocci, which are spherically shaped bacteria. The significance of a long axis or a cell pole measurement becomes unclear for such a morphology. Cytometry calculations must therefore be more generalized, or carefully checked, if analyzing morphologies which are not rod-shaped.

**TABLE S1.**
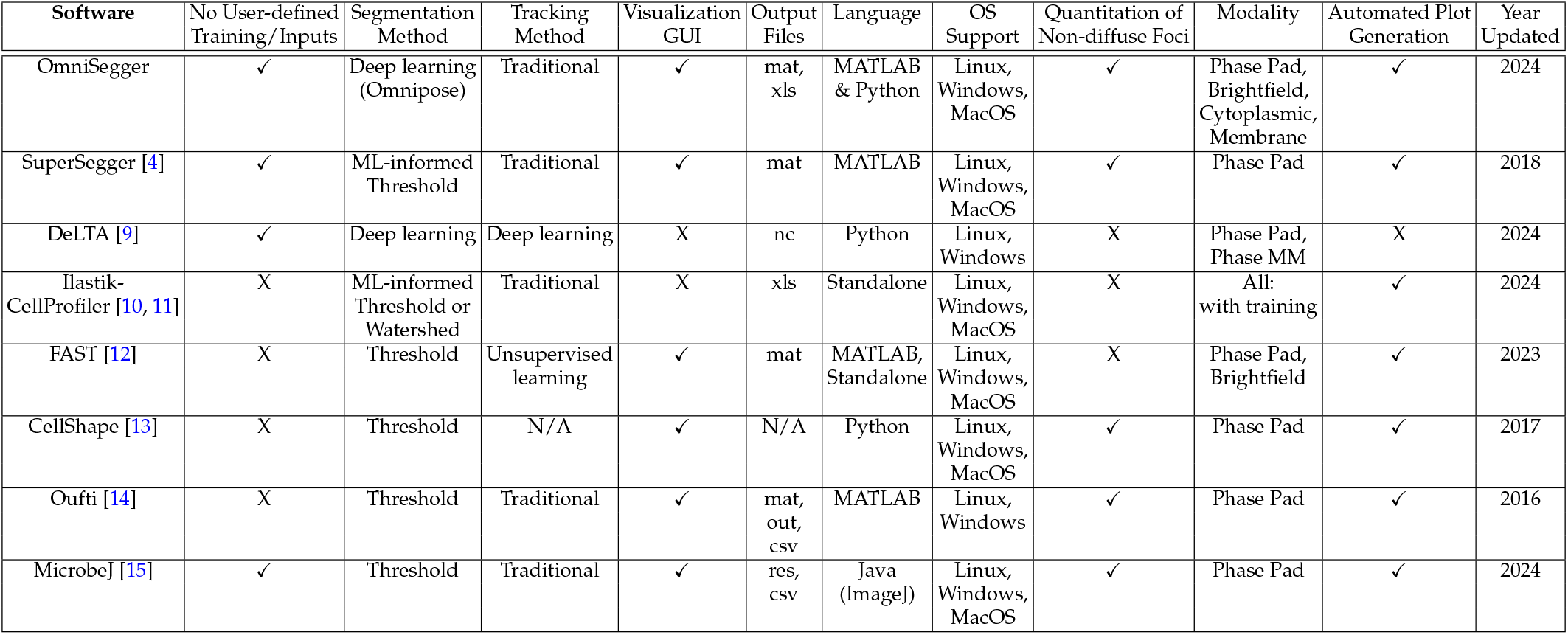
An extended comparison of features and functions for cellular imaging analysis software packages.

**FIG. S1.**
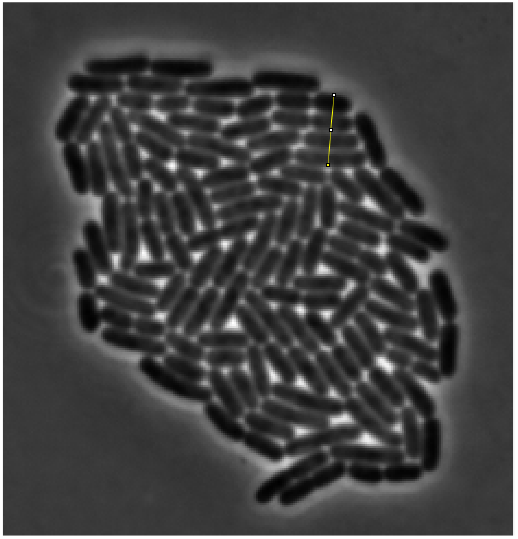
Cell length measurement across four cells from frame 138 of the time-lapse.

## Appendix C: Proliferation challenge: Cell width measurement

The cell width used as a proxy for sub-cellular accuracy was estimated in ImageJ by measuring the length of four parallel *E. coli* cells in contact (see Fig. S1). The length was then divided by four as an averaging method. We performed this measurement 4 times from frames 121, 133, 138, and 141 of the time-lapse. The average width is estimated to be 0.86 μm.

## Appendix D: Training the brightfield Omnipose model

We attempted to train a model to segment brightfield images taken in the focal plane of the bacterial samples. However, we noticed very poor performance of the model upon evaluation of in-focus test data (see Fig. S2).

After failing to train a model for images taken in the focal plane, we next attempted to train a model for segmentation of under- and over-focused images. The ground-truth dataset used to train the model does not include in-focus planes, only under- and over-focused planes in increments of 0.1 μm.

Because over-focused brightfield appears similar to phase-contrast, we generated approximate masks using the Omnipose phase-contrast model. Next, the masks were hand-annotated in Napari, informed by membrane-labeled fluorescence signal as validation when the fluorescence images were available, to refine and correct errors. Four to six colors were used to annotate the masks [3]. The masks were then converted to a standard 16-bit integer label mask PNG. The under- and over-focused images had a slight offset which were accounted for by performing a non-rigid image registration on the images with reference to the ground-truth masks.

In addition to our own images, the published bright-field images from DeepBacs [34] were included in the ground-truth dataset.

Number of images: 471.

Total cell count: 6623.

Further details about the datasets used for ground-truth annotation are included in Data Table S1.

## Appendix E: Materials and Methods

### 1. Bacterial sample preparation

#### Proliferation dataset

MG1655 *E. coli* was grown in Luria-Bertani media (LB) overnight, then reinoculated into fresh LB medium at a dilution of 1:1000, and grown to OD_600_ 0.1 before imaging. Cells were spotted onto a 4% agarose pad was prepared with M9 minimal media (1X M9 salts, 2 mM MgSO4, 0.1 mM CaCl2, 0.4% glycerol, and 0.2% casamino acids).

**FIG. S2.**
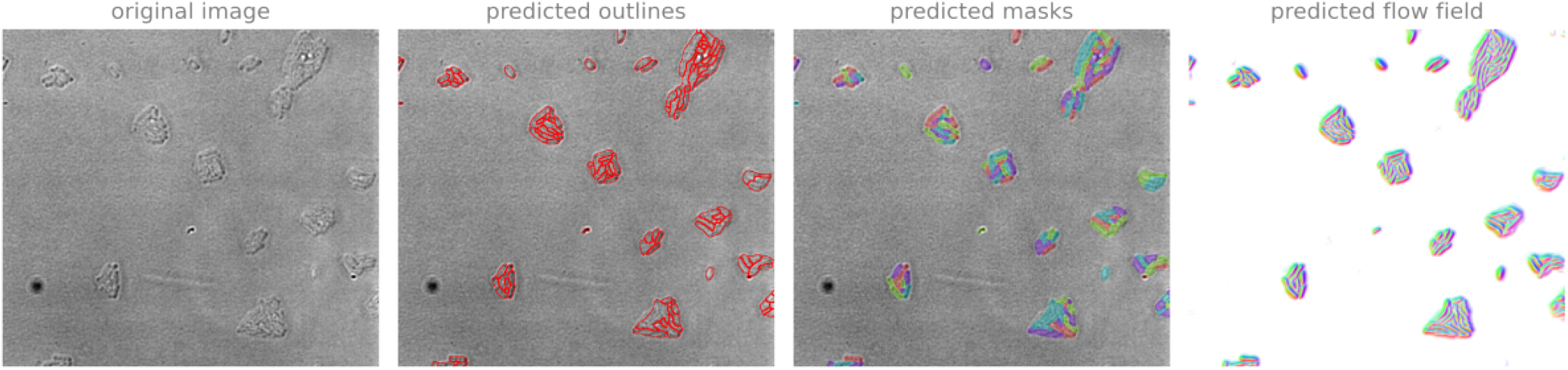
Brightfield model performance on in-focus image. Brightfield image of *E. coli* in the focal plane. The low contrast of in-focus brightfield makes the cells difficult to distinguish, both for the human eye and for segmentation algorithms.

#### Morphology dataset

MG1655 *E. coli* was grown in Luria-Bertani media (LB) overnight at 30°C, then reinoculated into fresh LB medium at a dilution of 1:1000 and grown for 1.5h at 30°C, then pelleted and resuspended in LB with 10μm hydroxyurea and grown for 1.5hr at 30°C. A 2% agarose pad was prepared using the same LB with 10μm hydroxyurea. Cells were spotted onto the pad and left in the microscope chamber at 37°C for 1hr before imaging.

#### Modality dataset-brightfield

TB28 attHK022 Plac::zipA-sfGFP bla pal-mCherry cat [35] was grown in Luria-Bertani media (LB) supplemented with 100μg/ml ampicillin overnight, then reinoculated into fresh M9 medium at a dilution of 1:100, and grown for 1h at 30°C before imaging. Cells were spotted onto a 2% agarose pad prepared with minimal M9 media (1X M9 salts, 2 mM MgSO4, 0.1 mM CaCl2, 0.2% glycerol, 10 μg/ml thiamine HCl) and left in the microscope chamber at 30°C for 1hr before imaging.

#### Modality dataset-cytoplasmic fluorescence

ASKA *lysC-GFP* (JW3984) was grown in a 96-deep well plate overnight in Luria-Bertani media (LB) supplemented with 34 μg/mL chloramphenicol (Cm34) at 30°C. The strain was then diluted 1:25 into M9 minimal media (1X M9 salts, 2 mM MgSO4, 0.1 mM CaCl2, 0.2% glycerol, 10 μg/ml thyamine HCl, and 0.2% casamino acids) with Cm34 and allowed to grow to mid-log. Prior to imaging, the fusion expression was induced with 500μm of Isopropyl *β*-d-1-thiogalactopyranoside (IPTG) for 40min. Cells were spotted onto a 2% agarose pad prepared with M9 media without IPTG [5].

#### Modality dataset-cytoplasmic fluorescence

ASKA *ygaW-GFP* (JW2645) was grown in a 96-deep well plate overnight in Luria-Bertani media (LB) supplemented with 34 μg/mL chloramphenicol (Cm34) at 30°C. The strain was then diluted 1:25 into M9 minimal media (1X M9 salts, 2 mM MgSO4, 0.1 mM CaCl2, 0.2% glycerol, 10 μg/ml thiamine HCl, and 0.2% casamino acids) with Cm34 and allowed to grow to mid-log. Prior to imaging, the fusion expression was induced with 50μm of Isopropyl *β*-d-1-thiogalactopyranoside (IPTG) for 40min. Cells were spotted onto a 2% agarose pad prepared with M9 media without IPTG [5].

### 2. Microscopy methods

Imaging was performed using a Nikon Eclipse Ti-E microscope, through a 60X 1.4 NA CFI oil-immersion Phase objective onto an Andor Neo sCMOS camera. The microscope chamber was heated to 30°C or 37°C.

### 3. Package versions

The following package versions were used to generate the data used in the figures:

- DeLTA: 2.0.5, main branch, commit 32e75d60; Python 3.11.10
- Ilastik 1.4.0, GPU-enabled
- CellProfiler 4
- SuperSegger: main branch, commit 6d58c6e; MATLAB R2024a
- OmniSegger: main branch, commit 22d9b69; MATLAB R2024a
  - Omnipose: main branch, commit a585929; Python 3.10.12

### 4. Computational resources

Analyses were performed with the following:

- OS: Ubuntu 24.04.1 LTS
- CPU: Intel(R) Core(TM) i9-9900K
- GPU: NVIDIA GeForce RTX 3090 Ti
- RAM: 32 GB

CellProfiler run on Windows 11 with Intel(R) Core(TM) i7-1165G7.

### 5. Image processing

The following commands/protocols were used for the analyses in this study:

- DeLTA - Command line `delta run -c 2D -i` `/dir/name/pos{p}cha{c}fra{t}.tif`
- Ilastik/CellProfiler: Followed protocols detailed on Youtube CellProfiler Tutorial: pixel-based classification with ilastik and at ImageProcessing-Benchmarking GitHub [21]
- SuperSegger - MATLAB `processExp(‘dir’)`
- OmniSegger - MATLAB `processExp(‘dir’)`

## Appendix F: Description of Supplementary Data

### 1. Data Table

**Data S1:** Bacterial strains and ground truth annotation for the Omnipose Brightfield model.

### 2. Datasets

**Data S2:** Proliferation dataset: phase-contrast images of wild-type MG1655 *E. coli* grown on a 4% agarose pad. Frame rate: 3min/frame. Raw images and OmniSegger masks.

**Data S3:** Morphology dataset: phase-contrast images of wild-type MG1655 *E. coli* treated with 10μm hydroxyurea growing on a 2% agarose pad. Frame rate: 5min/frame. Raw images and OmniSegger masks provided.

**Data S4:** Modality dataset-brightfield under-focused: under-focused brightfield images of TB28 attHK022 Plac::zipA-sfGFP bla pal-mCherry cat [35] growing on a 2% agarose pad. Images chosen from 0.5μm below the focal plane. Frame rate: 15min/frame. Raw images and OmniSegger masks provided.

**Data S5:** Modality dataset-brightfield over-focused: over-focused brightfield images of TB28 attHK022 Plac::zipA-sfGFP bla pal-mCherry cat [35] growing on a 2% agarose pad. Images chosen from 0.5μm above the focal plane. Frame rate: 15min/frame. Raw images and OmniSegger masks provided.

**Data S6:** Modality dataset-cytoplasmic fluorescence: fluorescence images of *lysC-GFP* fusion strain from the ASKA collection growing on a 2% agarose pad. Frame rate: 7min/frame. Raw images in both phase-contrast and fluorescence, and OmniSegger masks provided.

**Data S7:** Modality dataset-membrane fluorescence: fluorescence images of *cycA-GFP* fusion strain from the ASKA collection growing on a 2% agarose pad. Frame rate: 7min/frame. Raw images in both phase-contrast and fluorescence, and OmniSegger masks provided.

